# Going through phages: A Computational approach to Revealing the role of prophage in *Staphylococcus aureus*

**DOI:** 10.1101/2021.11.10.468171

**Authors:** Tyrome Sweet, Suzanne Sindi, Mark Sistrom

## Abstract

Prophages have important roles in virulence, antibiotic resistance and genome evolution in *Staphylococcus aureus*. Rapid growth in the number of sequenced *S. aureus* genomes allows for an investigation of prophage sequences in *S. aureus* at an unprecedented scale. We developed a computational pipeline to detect and analyze prophage sequences in nearly 10,011 *S. aureus* genomes, discovering thousands of putative prophage sequences with genes encoding virulence factors and antibiotic resistance.

**Importance:** Bacteriophages (phages) play key roles in bacterial evolution, governing abundance, adaptation and diversity of bacterial communities. Temperate phage can facilitate bacterial adaptation via transduction of novel genes. This study takes advantage of the unprecedented quantity of genomic sequencing in public repositories to analyze viral genes in 10,000 *Staphylococcus aureus* genomes. We found 196,727 prophage sequences, with an estimated total of 129,935 genes. We determined the function of these genes, identifying a large quantity of novel genes that benefit the host such as beta-lactamase, enterotoxins and cytotoxins. These results will inform studies of bacterial evolution and adaptation, by identifying the mechanism of horizontal transfer of genes that confer adaptive traits to bacteria, especially in the context of antibiotic resistance.

## Introduction

Bacteriophages are the most abundant self replicating organisms on earth, with an estimated global population of 10^31^, phages outnumber bacteria by 10 to 1^1^**(Liu, 2014)**. Lysogenic phage are transduced into the host bacterial genome as prophage sequences, and can have a range of selectional impacts on the host, spanning the breadth of the mutualism-parasitism continuum **(Blair, Webber,Baylay, Ogbolu and Piddock 2015)**. It is hypothesized that prophage sequences that confer a selective advantage to their host are more likely to be conserved in the bacterial genomes than those that are neutral or deleterious to their hosts **(Gandon 2016)**. The resultant expectation is that prophage sequences will contain an elevated quantity of genes conferring adaptive functions to host bacteria.

A well-studied example of an adaptive trait conferred by transduction by lysogenic phage is the mecA gene encoded in the phage Staphylococcus SCIURI7 **(Zeman, Mašlaňová, Indráková, Šiborová, Mikulášek, Bendíčková and Pantůček 2017)**. Transduction of this temperate phage into the Staphylococcus aureus genome confers resistance to broad spectrum beta-lactam antibiotics **(Scharn, Tenover and Goering 2013)**. Methicillin Resistant *Staphylococcus aureus* (MRSA) is one of the major causes of antibiotic resistant clinical infections. Between 1999 and 2005, hospitalizations for *S. aureus* increased from 294,570 patients to 477, 927. Moreover, MRSA was responsible for 127,036 patients in 1999 increasing to 278,203 by 2005 **(Klein, Smith and Laxminarayan 2007)**.

### Bacteriophages impact *S. aureus* evolution

Temperate bacteriophages, bacteriophages whose genome is incorporated into the host bacterium, can switch between the lytic and lysogenic life cycle **(Liu, 2014)**. The lytic cycle destroys the host, but as long as the phage stays lysogenic it provides several benefits. One benefit is protection from secondary phage attacks from other prophage. Temperate phages can lose their switching ability if there are mutations in the attachment sites. Changes to the gene that encode the recombinase responsible for the excision of phage can result in ‘grounding’ of the phage (**Ramisetty, and Sudhakari 2019**). Grounded phages offer the host benefits, without the risk of entering the lytic cycle.

*S. aureus* has a mesh-like cell wall composed of cross-linked polymer peptidoglycans (PG). Penicillin-binding proteins (PBPs), mediate the final stages of PG synthesis (**Scheffers and Pinho 2005**). Methicillin is a β-lactam antibiotic that inhibits the transpeptidation domain of PBPs, which weakens the cell wall (**Fishovitz, Hermoso, Chang and Mobashery 2014**). MRSA produces PBP2A due to the mecA gene that encodes it. Furthermore, this mecA gene is transducible by prophage (**Scharn, Tenover and Goering 2013**).

### Computational advances for Whole Genome Sequence (WGS) analysis

The number of sequenced and annotated phage genomes is relatively small with 40,981 phage sequences, and 266,129 prokaryotic genome sequences **(*Staphylococcus Aureus (ID 154) - Genome - NCBI*, n.d.)** on August 18th, 2018. A significant proportion of the genes encoded by both free living and prophage sequences are of unknown function (Touchon et al., 2016)**Moura, Criscuolo, Pouseele, Maury, Leclercq, Tarr, Brisse 2016)**. There is a large possibility for novel functions to be conferred to bacterial hosts by transduction by lysogenic phage **(Scharn, Tenover and Goering 2013)**. Given the exponential increase in the number of genome sequences deposited in public repositories, it is timely to take advantage of these sequences to analyze them for novel functions. In this study we analyze 10,011 *S. aureus* genomes downloaded from NCBI in 2018 for prophage sequences, and determine their functions. The total number of genome sequences for all organisms numbered 528,859 for 1 online repository**(*Genome List - Genome - NCBI*, n.d.)**. Advances in computational techniques for the analysis of large data sets have advanced the omics field by enabling researchers to analyze larger datasets at lower costs **(Krassowski, Das, Sahu, and Misra 2020)**.

In this study, we developed a computational pipeline to detect and analyze prophage sequences in nearly 10,000 *S. aureus* genomes. We discovered thousands of putative prophage sequences with genes encoding virulence factors and antibiotic resistance. In particular, we found genes encoding mecA, genes encoding toxins/antitoxins and clusters of prophage sequences that had genes in common. Our results, and methods developed, will facilitate similar studies for other bacterial species and promise to be a useful tool in the study of prophage host evolution.

## Materials and Methods

### *S. aureus* Genomes

*S. aureus* genomes were obtained from the National Center for Biotechnology Information NCBI’s Genbank repository on August 18, 2018 **(*Staphylococcus Aureus (ID 154) - Genome - NCBI*, n.d.)**. All available genome sequences (n=10,011 including complete and partial assemblies) were downloaded for this study. (Accession numbers are provided in Supplementary Tables 1&2).

### Viral Detection

Putative prophage sequences were detected using PhiSpy, Version 3.2 **(Akhter, Aziz and Edwards 2012)**. PhiSpy uses a random forest algorithm that has been trained on seven distinct features of prophage: protein length, transcription strand directionality, AT and GC skew, the abundance of unique phage words, phage insertion points and the similarity of phage proteins. PhiSpy has 49 available training sets to increase accuracy for specific genomes. We used the S. *aureus* training dataset (option 24) and identified 196,727 phage regions in our 10,000 *S. aureus* genomes.

### Prophage Clustering

Prophage sequences identified by PhiSpy were unique within a genome, but highly redundant between genomes. We identified highly-similar prophages between genomes through a reciprocal blast **(Johnson, Zaretskaya, Raytselis, Merezhuk, McGinnis and Madden 2008)** search. We increased the max_target_seqs to 12,000 (higher than our total number of *S. aureus* genomes) to ensure we captured all possible matches. We also used a custom output format which provided additional information on the alignment.

We then grouped prophages by using an undirected graph approach with nodes of the form: Genome *i*, Prophage *j*. Edges were added between nodes if they had a blast alignment which exceeded 90% similarity and 90% coverage of both source and target. We then identified genomes sharing the same prophage by determining the connected components, resulting in 191 unique phage clusters.

### Cluster Validation

Each of the 191 phage clusters were aligned with Muscle v3.8.1551 **(Edgar, 2004)** and ClustalW v2.1 **(Thompson et al., 1994)** to ensure each phage was similar. A score of 0.0000 indicates that the undirected graph script formed accurate phage clusters.

### Genome Annotation

One representative was selected from each of the 191 phage clusters and analyzed with 2 different tools for gene annotation: VGAS **(Zhang et al., 2019)**, and Prokka **(Seemann 2014)**. VGAS and PROKKA identified ORFS in each of the phage genome sequences. VGAS identifies ORFs through an enhanced version of the ZCurve algorithm **(Guo et al., 2003)**that was customized by adding 13 additional identifying variables (45 total) for the classification model, and BLASTP **(Mahram & Herbordt, 2015)**searches for gene prediction. The all ORFs were annotated by all both tools with default settings.

### Heat Maps

We identified shared genes between phage through a reciprocal blast search using the annotated phage sequences. We constructed a new undirected graph with the nodes being the phage genome and the edges representing genes shared between the phage. The output was a .csv file that listed each of the 191 phage with the genes shared with other phages. A pairwise count matrix was formed with the x and y axis’ are the 191 phage, and the cells show the number of genes shared. Using this matrix, a heatmap is created using the seaborn python package **(Bisong 2019)**. **(See Figures 4 and 7 in the results section)**

### Network Visualization

We used the layout_with_mds option for the layout function of the R package Igraph **(Csardi and Nepusz 2006)** to visualize the phages with shared genes using the pairwise count matrix for both PROKKA and VGAS.

### Jaccard Index

The Jaccard Index **(Hamers 1989)** was calculated using a modified version of the Jaccard index function in R **(Woudenberg 2021)** to compare the Prokka and VGAS networks.

## Results

Of the 10,011 genomes initially analyzed, there were 12 that were not annotated completely and did not pass the conversion to SEED **(Aziz, Devoid, Disz, Edwards, Henry, Olsen and Xia 2012)** due to missing locus tags **(*Prokaryotic Genome Annotation Guide*, n.d.)**. A further four genome sequences were too short, and did not have enough base pairs for PhiSpy to detect phage regions, resulting in a total of 9,995 genomes which passed initial QC and were used for subsequent analysis. Within these, we detected a total of 196,727 prophage sequences across the 10,011 genomes, with an average of 39 (standard deviation = 1967.468) prophage sequences per genome **(Figure 2)**.

**Figure 1:**
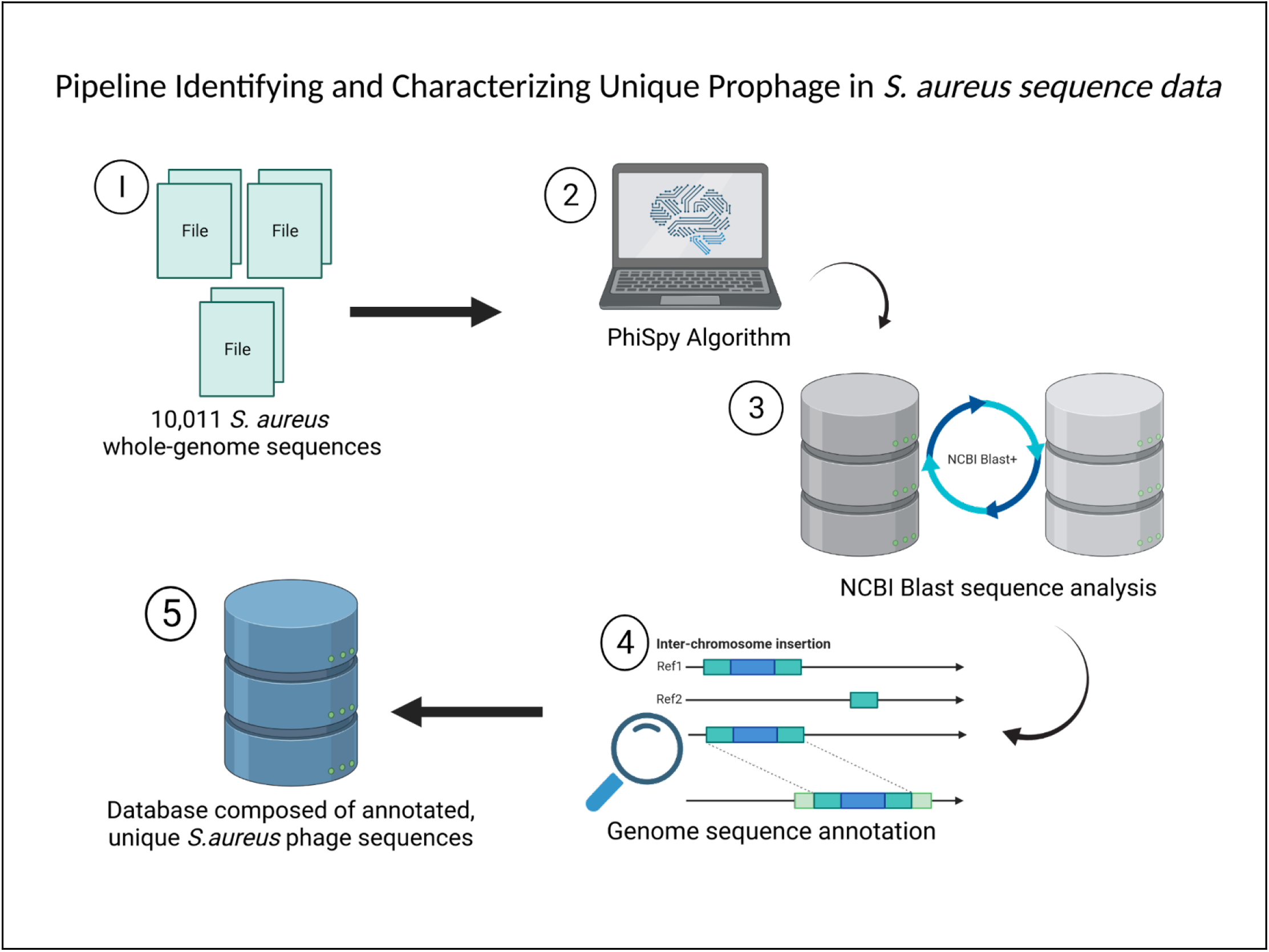
Pipeline Identifying and Characterizing Unique Prophage in *S. aureus* sequence data. A visualization of the workflow used to identify unique prophage sequences. **1)** 10,011 *S. aureus* genome sequences were downloaded from the National Center for Biotechnology information (NCBI). **2)** The sequences were analyzed by PhiSpy. **3)** The fasta files for each predicted prophage were compared against each other using NCBI Blast nucleotide alignment tool. Prophage sequences that had 90% similarity along their full length were considered to be the same. **4)** Phage sequences were annotated using two independent methods (VGAS, Prokka) e. **5)** The resulting database of annotated, unique phage sequence allows for the identification of gene function encoded within prophage in *S. aureus.* (See **materials and methods** section above for more information)

**Figure 2:**
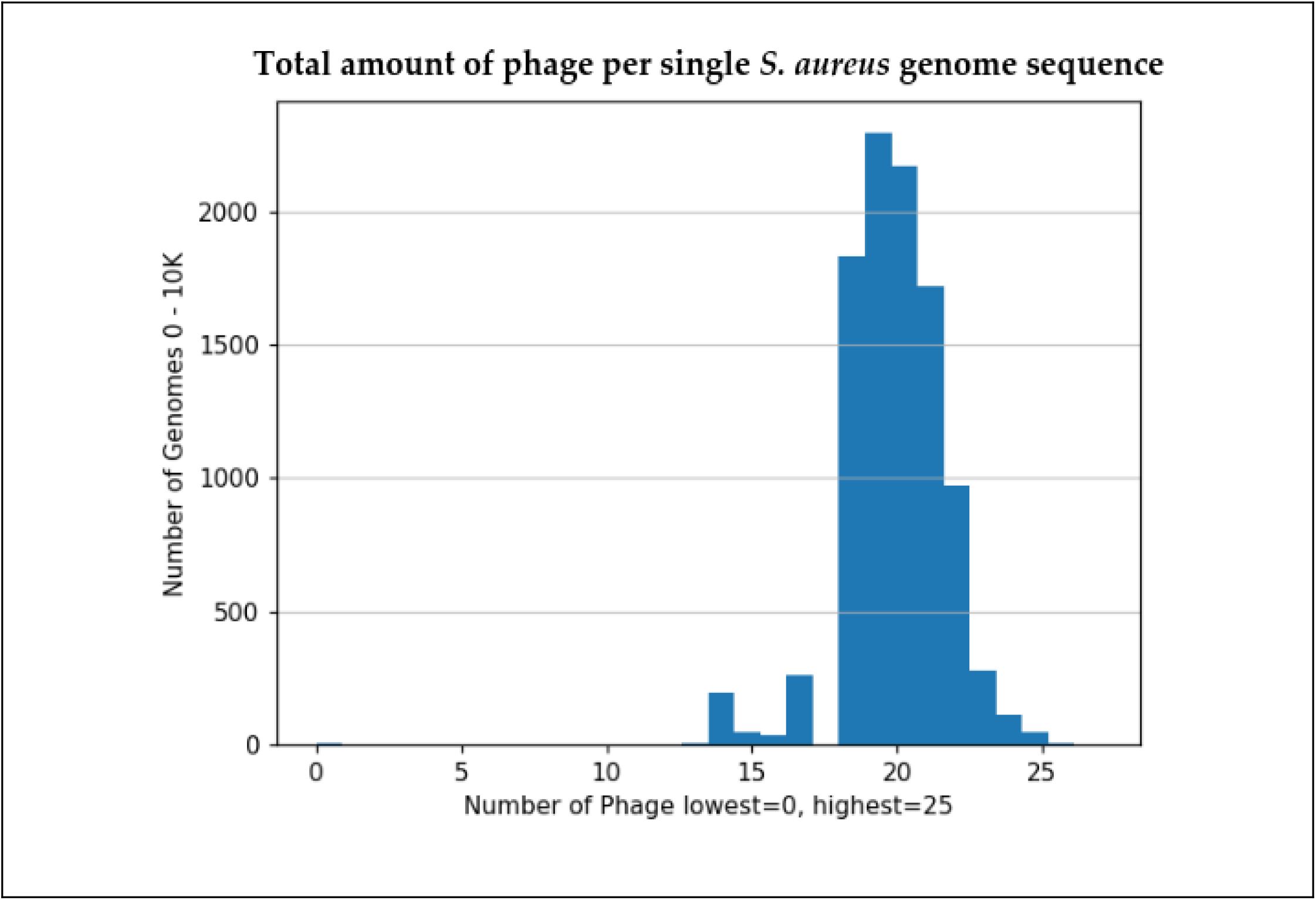
Total amount of phage per single *S. aureus* genome sequence. Detected a total of 196,727 prophage sequences across the 10,000 genomes. There is an average of 39 (standard deviation = 1967.468) prophage sequences per genome. 1125 *S. aureus* sequences had 25 phage regions present, and 4 genomes had 0 phage detected. The x-axis reflects the exact totals each genome contained per genome (y-axis). (see **Analysis Uncovers 191 Unique Prophage Sequences** section above for more information).

### Analysis Uncovers 191 Unique Prophage Sequences

Reciprocal BLAST analysis coupled with undirected graph analysis (see Methods) found that the 196,727 prophage sequences corresponded to 191 unique prophage sequences. Each unique prophage sequence appeared in an average of 1024 host genomes (Standard deviation = 2581.33)**(Figure 3)**. Each prophage contained an average of 16.83 putative coding regions, resulting in a total of 3,208 (VGAS) and 3,203 (Prokka) unique open reading frames (ORFs). One phage appeared in all 9,995 genome sequences, while 42 different phages were found in only a single genome sequence.

**Figure 3:**
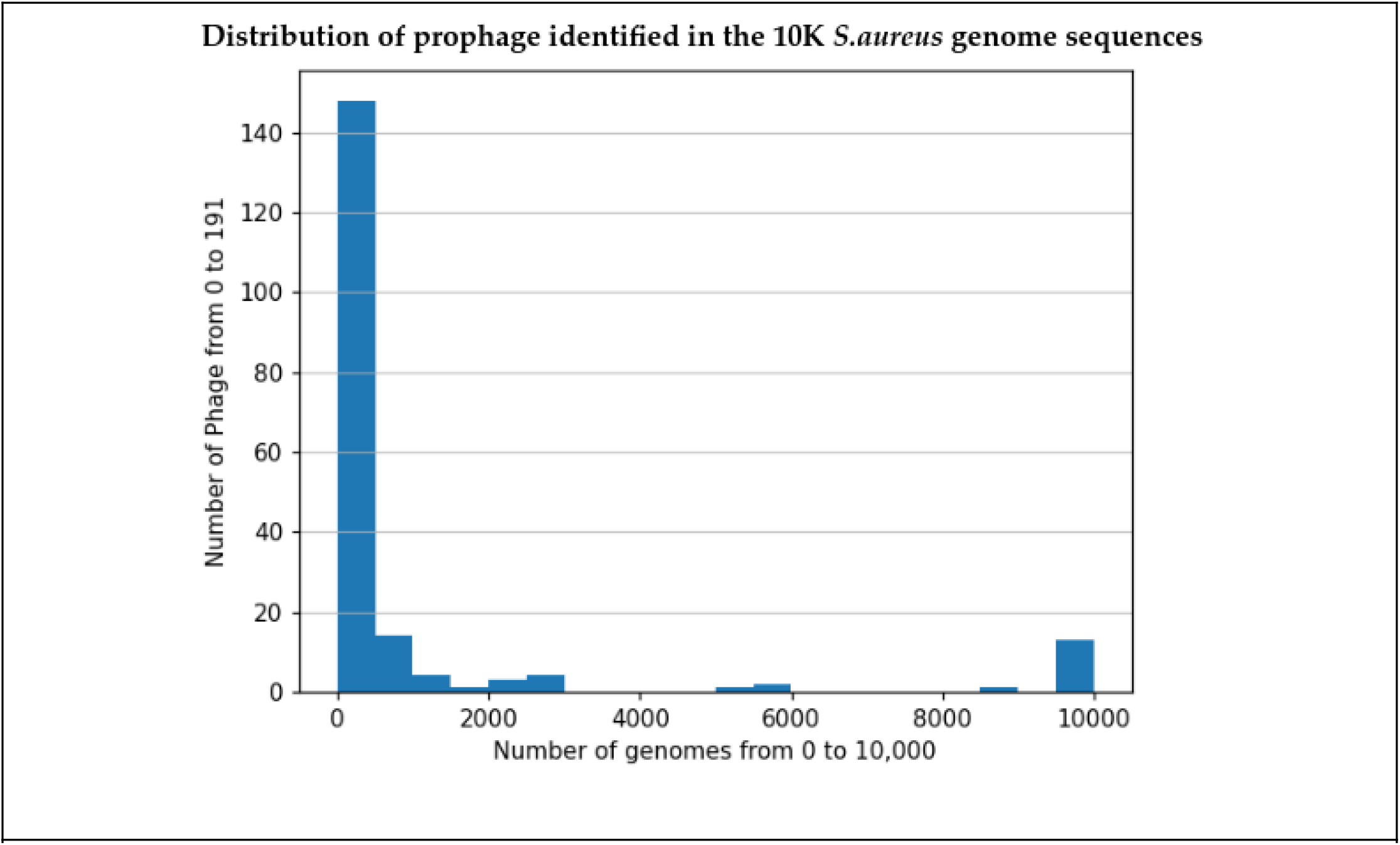
Distribution of prophage identified in the 10K *S.aureus* genome sequences. There were a few phage sequences detected in nearly every genome sequence. 191 prophages with an average of 1024 (Standard deviation = 2581.33). The highest number of genomes a single phage was detected is 9995, and the lowest was 1. (see **Analysis Uncovers 191 Unique Prophage Sequences** section above for more information).

### Analysis Detects Thousands of ORFs with Potential Gene Function

One representative was selected from each of the 191 phage clusters and analyzed with two different tools for gene annotation: VGAS **(Zhang et al., 2019)**, and Prokka **(Seemann, 2014)**. VGAS identified 3,208 genes in the 191 unique phage, and PROKKA detected 3,203 genes. For the PROKKA results, 1,155 ORFs did not have an identified function. 807 predicted ORFs corresponded to known ORFs with accession numbers matching known databases ISfinder **(Siguier, Pérochon, Lestrade, Mahillon and Chandler 2006)**, NCBI **(*National Database of Antibiotic Resistant Organisms (NDARO) - Pathogen Detection - NCBI*, n.d.)**, UniProtKB **(Boutet, Lieberherr, Tognolli, Schneider and Bairoch 2007)**. 2041 genes had a predicted gene function. VGAS predicted 2935 ORFs, 362 of which corresponded to known accession numbers matching databases Swissprott and refseq **(Pruitt and Maglott 2001)** and 309 other predicted ORFs had predicted gene functions.

### Analysis Shows Shared ORFs between Unique Prophage Sequences

2 undirected graphs based on the genes identified by PROKKA and VGAS were created. The approach outlined in the **“prophage clustering”** section with nodes of the form: Genome *i*, identified gene *j*. Edges were added between nodes if they had a matching identified gene. We then Compared the edges produced by both tools PROKKA and VGAS with each other.

We found a total of 1,335 shared edges defined by PROKKA and VGAS. The lowest number of shared edges between phage sequences was 1, and the highest was 73 (**Figure 4A**). There were 1,306 shared edges between PROKKA and VGAS, and 28 shared edges unique to PROKKA (**Figure 6**) out of the total 1,335 (**Figure 5**). In the 28 unique PROKKA the numbers of shared edges between each node ranged from 1 to 22. VGAS defined a total of 1,334 connected components. The lowest amount of genes shared between phage sequences was 1, and the highest was 75 (**Figure 4**). There were 27 shared edges unique to VGAS (**Figure 9**) out of the total 1334 (**Figure 8**). The 27 unique VGAS shared edges ranged from 1 to 22 as well.

**Figure 4:**
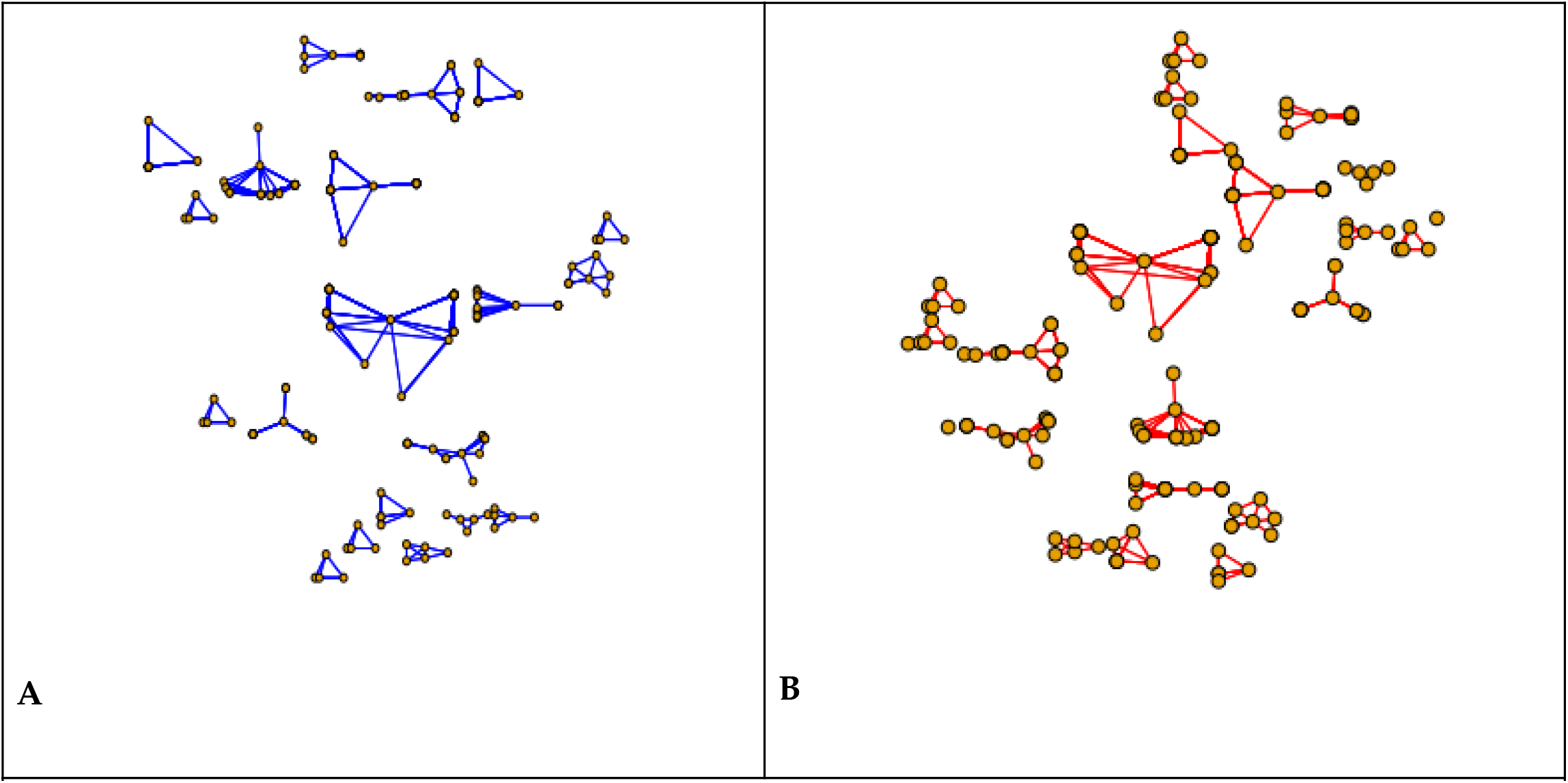
PROKKA and VGAS Undirected Graphs shows shared ORFs between unique prophage sequences. **A)** This graph shows the relationship between phage genomes by their gene content. Specifically, the nodes represent the 191 phage genomes and edges between nodes indicate the two phage share a gene (as annotated by Prokka). We determined that there were 1335 connected components with shared genes ranging from size 1 to 73. **B)** This graph shows the relationship between phage genomes by their gene content. Specifically, the nodes represent the 191 phage genomes and edges between nodes indicate the two phage share a gene (as annotated by VGAS). We determined that there were 1334 connected components with shared genes ranging from size 1 to 75. (See **Analysis Shows Shared ORFs between Unique Prophage Sequences** section above for more information).

**Figure 5:**
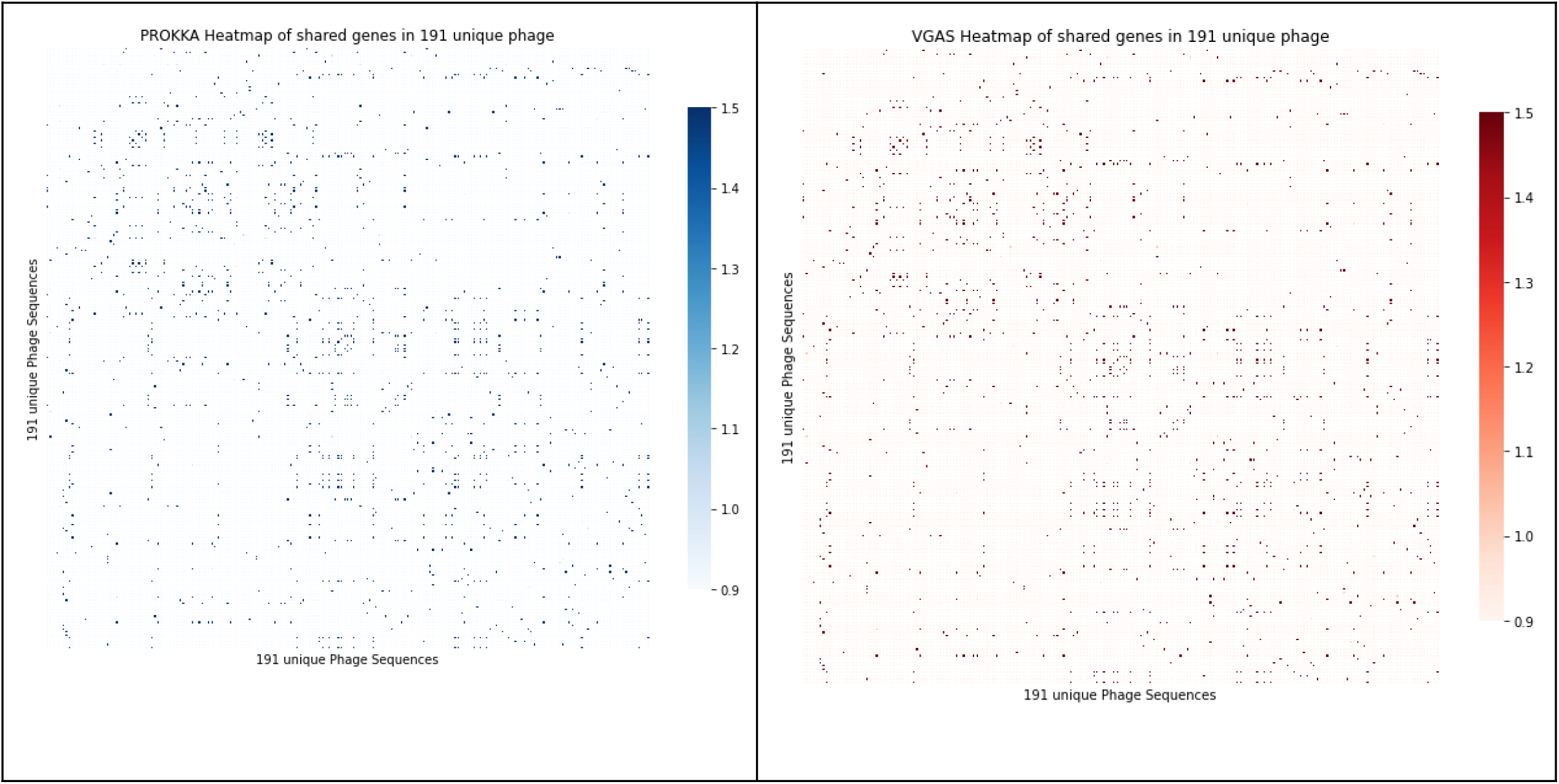
Heatmaps showing distribution of shared orfs between 191 unique phage sequences. **Left)** This heatmap shows the numbers of genes shared between phage genomes as annotated by Prokka. These numbers ranged from 1 to 73. The X and Y axis are the 191 unique phage sequences. **Right)** This heatmap shows the numbers of genes shared between phage genomes as annotated by VGAS. These numbers ranged from 1 to 75. The X and Y axis are the 191 unique phage sequences. ( See **Heat Maps** section above for more information).

**Figure 6:**
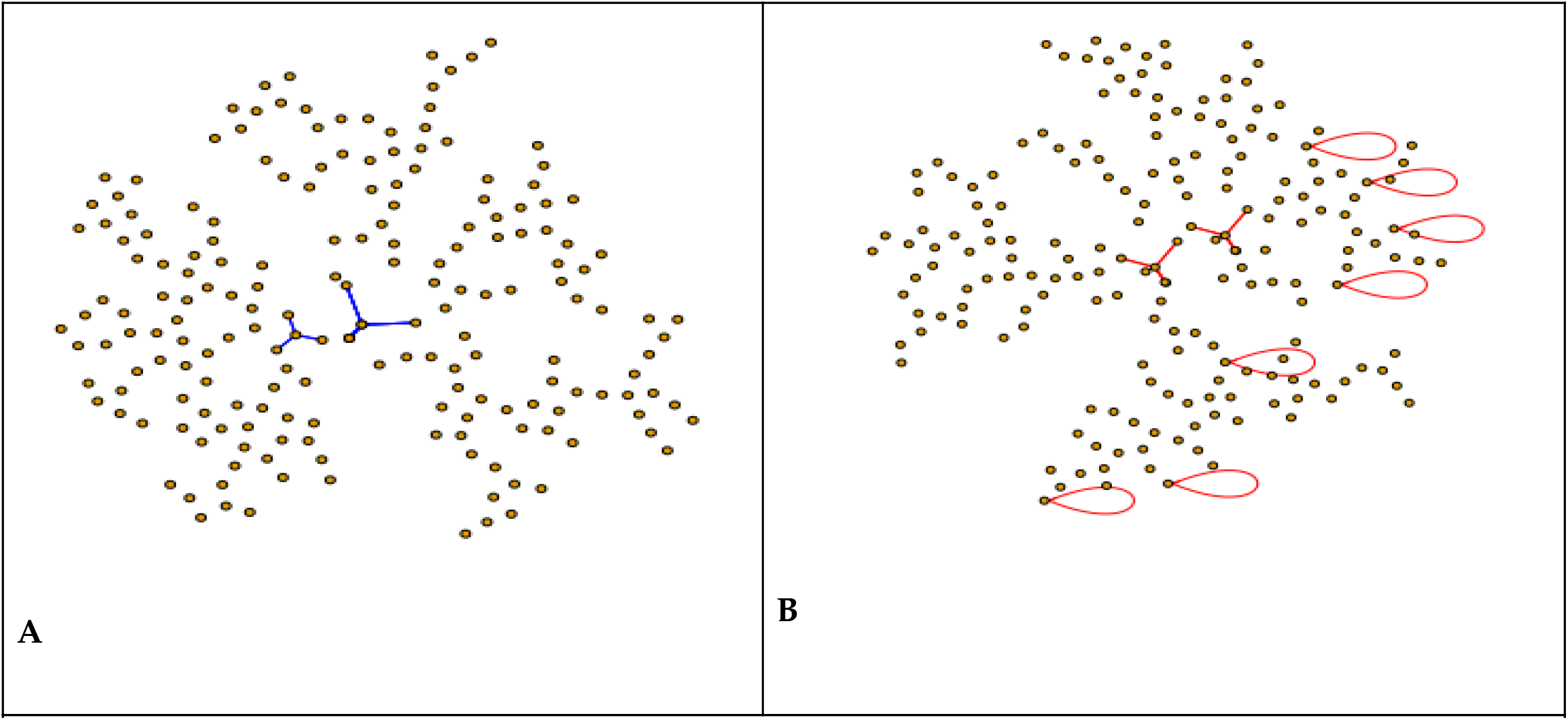
PROKKA and VGAS Undirected graph shows differences in shared ORFs between unique prophage sequences. **A)** 28 unique edges, 1306 shared with VGAS **B)** 27 unique edges, 1306 shared with Prokka. Both the PROKKA and VGAS graphs shared the same range of connections with 1 as the lowest, and the highest at 22. (See **Analysis Shows Shared ORFs between Unique Prophage Sequences** section above for more information).

## Discussion

There were several virulence factors and toxins identified in the 191 unique prophage representatives, 1% of the total 196,727 phage detected. ***Staphylococcus aureus subsp. aureus strain NCTC 8325*** is referenced several times throughout the dataset. It was used as a propagating strain for bacteriophage 47 of the international typing set of bacteriophages and is considered the prototypical strain for most genetic research on *S. aureus* (Staphylococcus Aureus Subsp. Aureus NCTC 8325 (ID 57795) - BioProject - NCBI, n.d.).

### Genes Encoding mecA Found in 2 of the 191 Unique Prophage

There were several traces of antimicrobial resistance found in the 191 phage clusters. The *mecA* ancestral gene specifically was identified in 2 sequences. The first sequence, accession number ASM900v1**(*Staphylococcus Aureus (ID 154) - Genome - NCBI*, n.d.)**, cluster group has 1023 phage, 10% of the total *S. aureus* genomes. ASM900v1, or RF122 (ET3-1) provides a framework for the identification of specific factors associated with host specificity in this major human and animal pathogen **(Herron et al., 2002)**. RF122 (ET3-1) has several genes involved in host colonization, toxin production, iron metabolism, antibiotic resistance, and gene regulation **(Herron-Olson et al, 2007)**. ASM323779v1 **(*Staphylococcus Aureus (ID 400143) - BioProject - NCBI*, n.d.)** is the only phage in the cluster, making it individually unique compared to the 196,727 total detected. It is a part of 184 S. aureus isolates collected from 135 patients over a timespan of 3 years at an Italian paediatric hospital **(Manara et al., 2018)**.

### 48 Unique Gene Functions Supporting Antimicrobial Resistance

48 unique encoding traces of Antimicrobial Resistance **(Table 1)**. 4 genes stuck out the most due to the number of clusters they appeared in. GDAEFEPF_00005 Staphylococcal complement inhibitor, a gene found in ASM2514v1 appeared in 10 clusters **(Nübel et al., 2010)**. GHDFECEE_00007 Superantigen-like protein 13 was found in ASM17451v1 and appeared in 8 clusters **(Cameron et al., 2012)**. ASM17451 also contained GHDFECEE_00008 Superantigen-like protein 13 which appeared in 7 clusters. GAIDFPLK_00004 Superantigen-like protein 13 was found in ASM1150v1 and was identified in 7 clusters **(Holden et al., 2004)**.

### 4 Genes Showing Traces of Toxin/Antitoxin (TA) System

Toxin/Antitoxin (TA) systems encode toxin proteins that interfere with vital cellular functions and are counteracted by antitoxins. There are 6 different types of TA systems, S. aureus has genes identified showing types I, II and III **(Schuster & Bertram, 2016)**. Type I toxin-antitoxin systems have the base-pairing of antitoxin RNA with the toxin mRNA **(Fozo et al., 2008)**. Type III systems toxic proteins and an RNA antitoxin have a direct iteration where the toxic proteins are neutralized by the RNA gene **(Labrie et al., 2010)**.

Type II, the most studied TA system, has proteic antitoxin that tightly binds and inhibits the activity of a stable toxin **(Hayes, 2003)**. The TA system yoeB-yefM has been detected as genes MBJHDCJA_00021 Toxin YoeB and MBJHDCJA_00022 Antitoxin YefM in ASM900v1 **(Herron et al., 2002; Herron-Olson et al., 2007)**. yoeB inhibits bacterial growth and translation by cleavage of mRNA molecules and is repressed by antitoxin yefM **(Schuster & Bertram, 2016)**. Enterotoxin Type A causes food poisoning and was identified in 3 genome sequences **(Ono et al., 2012)**. M1022 (NCTC 8325) was identified in 2 genome sequences (Staphylococcus Aureus Subsp. Aureus NCTC 8325 (ID 57795) - BioProject - NCBI, n.d.). CAFLMJIC_00063 Enterotoxin type A was identified in 1 genome sequence **(Herron et al., 2002; Herron-Olson et al., 2007)**. **(See Tables 1 and 2 the supplemental materials section)**

### 13 Most Shared Genes in the 191 Unique Phage

4 genes that stand out the most due to the amount of phage they were found in **(table 2)**. KHDAMHGJ_00009 Chorismate synthase, found in M0471 (*Staphylococcus Aureus Subsp. Aureus NCTC 8325 (ID 57795) - BioProject - NCBI*, n.d.), was identified in 17 phage clusters. Its gene function is shikimate pathway, which shows signs of AMR in plants **(Mander and Liu 2010)**. EOLKNJBM_00007 Nucleoside diphosphate kinase in ASM1150v1_genomic.gbff_pp18.ffn **(Holden et al., 2004)** was found in 16 phage clusters. MIIMDJNA_00002 Heptaprenyl diphosphate synthase component 2 in ASM24879 (*Staphylococcus Aureus Subsp. Aureus CIG1612 (ID 60683) - BioProject - NCBI*, n.d.) was identified in 15 clusters. HGDEFLKI_00006 3-dehydroquinate synthase in M0877_V1_genomic.gbff_pp18.ffn (*Staphylococcus Aureus Subsp. Aureus NCTC 8325 (ID 57795) - BioProject - NCBI*, n.d.) was identified in 14 phage clusters.

## Acknowledgments

- Multi-Environment Computer for Exploration and Discovery (MERCED) cluster at UC Merced, funded by National Science Foundation Grant No. ACI-1429783
- National Science Foundation –National Research Traineeship in Intelligent Adaptive Systems (NRT-IAS) (Award No. 1633722)
- National GEM Consortium/Georgia Tech Research Institute - GEM PhD Engineering and Science Fellowship
- Ali Heydari

